# Gustatory-neuron-supplied R-spondin-2 is required for taste bud replenishment

**DOI:** 10.1101/2024.02.21.581408

**Authors:** Jiang Xu, Alan Moreira de Araujo, Ranhui Xi, Xiaoli Lin, Chanyi Lu, Minliang Zhou, Kurt Hankenson, Robert F. Margolskee, Ichiro Matsumoto, Guillaume de Lartigue, Myunghwan Choi, Peihua Jiang

**Author notes:** Equal contributions (Jiang Xu, Alan Moreira de Araujo). Corresponding author: Peihua Jiang, Monell Chemical Senses Center, 3500 Market Street, Philadelphia, PA 19104.

## Abstract

Taste buds undergo continuous cell turnover throughout life, and taste cell replenishment relies strictly on innervation, a phenomenon first described almost 150 years ago. Recently, we provided evidence that R-spondin 2 (Rspo2) may be the long-sought gustatory neuron-supplied factor that regulates taste stem cell activity, via its interaction with taste stem/progenitor cell-expressed receptor Rnf43/Znrf3. Yet, whether gustatory-neuron-supplied Rspo2 is strictly required for taste tissue maintenance has not been resolved. Here, we set out to determine the necessity of gustatory-neuron-supplied Rspo2 in taste tissue homeostasis using genetic approaches. We used a mouse line that harbors the neomycin-resistance gene (*NeoR*) in one of the intron regions of the *Rspo2* gene, which results in reduced expression of Rspo2. The number of taste buds is significantly reduced in these mice, compared to wild-type mice, in both anterior and posterior tongue. This phenotypic change was completely reversed by removing *NeoR* from the *Rspo2* gene, thus making it normal. We also combined adeno-associated virus (AAV)-based delivery of Cre recombinase with a mouse line amenable to Cre-based ablation of the *Rspo2* exons encoding the receptor-binding domains. Such deletion of Rspo2 in the nodose-petrosal-jugular ganglion complex led to nearly complete loss of taste buds in the circumvallate papilla. Thus, we demonstrate that Rspo2 is the long-sought gustatory-neuron-supplied factor that acts on taste stem cells to maintain taste tissue homeostasis.

**Significance:** We have known for 150 years that innervation is required to induce and maintain cell replacement in taste buds. Until recently, the identity of the inducing factor produced by neurons was unknown. We have shown that R-spondin alone is sufficient to substitute for neuronal input to induce taste bud regeneration. Using a genetic loss-of-function approach, we now demonstrate that gustatory-neuron-expressed Rspo2 is required to maintain taste tissue homeostasis. Altogether, our work reveals that Rspo2 is the long-sought neuron-supplied factor that regulates the activity of taste stem/progenitor cells.

## INTRODUCTION

Humans and other species rely on the sense of taste to evaluate food prior to ingestion (1–4). For instance, sweet and umami tastes indicate calorie-rich, nutritious food items and trigger ingestion, whereas bitter taste in food indicates potentially harmful and toxic substances and triggers avoidance (1–4). Taste buds, strategically scattered throughout the oral cavity and upper alimentary tract (i.e., epiglottis), are peripheral sensory organs that detect taste stimuli in food and drink. Each taste bud, organized in an onion-shaped structure, comprises about 50-100 cells, including taste receptor cells that detect taste stimuli and supporting cells that do not. Taste buds are innervated by gustatory neurons located in a few cranial nerve ganglia [e.g., geniculate ganglion and nodose-petrosal-jugular ganglion (NPJ) complex], which relay taste information to the brain to register taste sensation/perception (1–4).

Like other epithelial cells, taste bud cells turn over constantly throughout life (5, 6). To maintain taste tissue homeostasis, adult taste stem/progenitor cells (e.g., cells positive for Lgr5 in posterior tongue) continue to generate new cells to replace senescent ones (7–10). Unlike in other epithelial tissues, in taste buds maintenance and regeneration are strictly dependent on innervation, which was noted almost 150 years ago (11, 12).

Yet, single isolated Lgr5^+^ or Lgr6^+^ taste stem/progenitor cells can grow into ever-expanding 3-D structures (termed as “taste organoids”) and generate taste cells in the absence of neurons or other types of cells *ex vivo*, suggesting that a soluble factor in the organoid culture medium may substitute for neuronal input to activate taste stem/progenitor cells to produce taste cells (13–16). As in other organoid culture systems, R-spondin, EGF, and Noggin are key components of the medium used for culturing taste organoids (17, 18).

In pursuit of the gustatory-neuron-supplied factor that regulates taste tissue homeostasis, we recently found that R-spondin-2 (Rspo2; one of the four vertebrate R-spondin proteins) is abundantly expressed in gustatory neurons and demonstrated that exogenous R-spondin (e.g., Rspo1 or Rspo2) is sufficient to promote generation of taste buds after denervation (19). Furthermore, we recently showed that epithelial-cell-specific ablation of two E3 ubiquitin ligases, Rnf43/Znrf3, that are receptors for R-spondin (binding of R-spondin to Rnf43/Znrf3 silences their ligase activity) reproduces (phenocopies) the effect of R-spondin on taste tissue homeostasis, such as rendering innervation dispensable for taste bud maintenance and regeneration (20). Although our previous work demonstrates the *sufficiency* of Rspo2 in maintaining taste tissue homeostasis, the *necessity* of Rspo2 remained unclear. Here, we used genetically engineered mice to determine whether gustatory-neuron-supplied Rspo2 is necessary to maintain taste bud structure.

## RESULTS

### The number of taste buds in the circumvallate papilla is greatly reduced in genetically engineered *Rspo2^Neo/Neo^* mice, with a gene-dosage-dependent effect

To determine the necessity of gustatory-neuron-supplied Rspo2 for maintaining taste tissue homeostasis, we used a genetically engineered mouse strain, termed *Rspo2^Neo/Neo^*, in which two LoxP sites along with the sequence encoding the neomycin resistance gene cassette (*NeoR*) were inserted into the intron regions that flank exons 4 and 5 of the *Rspo2* gene (**Fig. 1A**). This mouse line enables removal of the Lgr4/5/6 binding domain in a Cre recombinase-dependent manner without affecting other domains (e.g., the Rnf43/Znrf3 binding domain encoded by exon 3), because coding exons of *Rspo2* are all in-frame (**Fig. 1A**) (21). Initially, we crossed this strain with the *Thy1-CreERT2, -EYFP* (SLICK-H [single-neuron labeling with inducible Cre-mediated knock-out]) strain (22), in which the transgene was known to be widely expressed throughout the peripheral and central nervous system, to enable inducible deletion of a floxed allele in the transgene-expressing neurons. Using this approach, we tested if tamoxifen-induced ablation of the Lgr4/5/6 binding domain of *Rspo2* would lead to changes in taste structures. Surprisingly, we found a pronounced reduction in the number of taste buds in *Rspo2^Neo/Neo^*homozygous mice regardless of the presence or absence of the *Thy1-CreERT2, -EYFP* driver (**Fig. S1**), suggesting this phenotype is constitutive, not induced by Thy1-CreERT2. Furthermore, when we examined *Thy1-CreERT2, -EYFP* mice for intrinsic YFP in the NPJ complex, only a small subset of neurons were YFP positive, arguing against the effectiveness of using this *Thy1-CreERT2* strain as a Cre driver to knock out genes in gustatory neurons (**Fig. S2**).

**Fig. 1.**
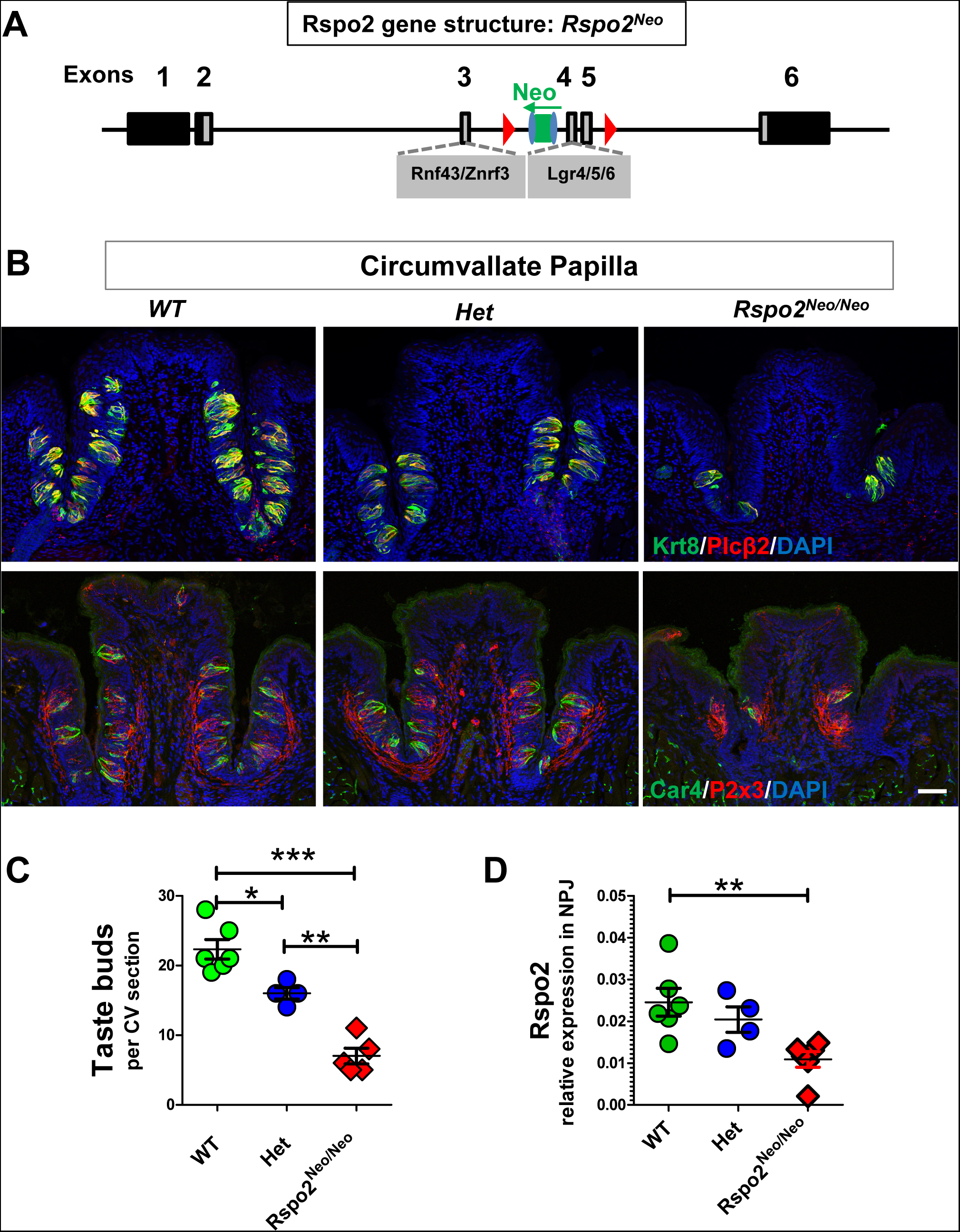
Fewer taste buds in the circumvallate papilla of mice harboring a hypomorphic allele of *Rspo2* than in wild-type mice. **A)** Schematic drawing illustrating the genetic engineering of the *Rspo2* locus in mice carrying the *Rspo2^Neo^* allele. Red triangles depict LoxP sites. The *NeoR* gene cassette includes two FRT sites (blue ovals) and the *NeoR* gene (green rectangle). **B)** Representative confocal images of circumvallate papilla sections of wild-type (WT), heterozygous (*Rspo2^Neo/+^*; Het), and homozygous *Rspo2^Neo/Neo^*mice, immunostained with antibodies against Krt8 (green) and Plcβ2 (red; top row) or against Car4 (green) and P2x3 (red; bottom row). Images are compressed z-stacks and were acquired using the same settings for each antibody. Scale bar: 50 μm. **C)** Quantification of taste buds that comprise Krt8+ and Plcβ2+ cells in the circumvallate papilla of wild-type (WT), heterozygous (Het), and *Rspo2^Neo/Neo^* mice. Significant difference was noted among different genotype groups (* p <0.05; ** p <0.01; *** p <0.001). Each point represents a single mouse. Four to six mice were used for each genotype. **D)** qRT-PCR analysis of the expression of Rspo2 in NPJ of wild-type (WT), heterozygous (Het), and *Rspo2^Neo/Neo^* mice. Significant difference was noted in the amount of NPJ-expressed Rspo2 between wild-type and *Rspo2^Neo/Neo^* mice (** p <0.01). Each point represents data generated using two NPJ (combination of both left and right) from a single mouse. Four to six mice were analyzed for each genotype.

To avoid any potential complication from the *Thy1-CreERT2, -EYFP* crossing, we directly examined three strains: homozygous *Rspo2^Neo/Neo^*, heterozygous (*Rspo2^Neo/+^*), and wild-type control mice (not crossed with *Thy1-CreERT2, -EYFP*). We performed double immunostaining of circumvallate papilla sections using antibodies against Krt8 (a general taste bud cell marker) and Plcβ2 (a type II cell marker) (**Fig 1B, top**) or against Car4 (a type III cell marker) and P2x3 (a gustatory nerve marker) (**Fig. 1B, bottom**). The morphology of existing taste buds in *Rspo2^Neo/Neo^*mice appeared to be normal and innervated. However, compared to wild-type and heterozygous mice, *Rspo2^Neo/Neo^* mice showed a significant reduction in the number of taste buds in the circumvallate papilla, as expected (**Fig. 1B,C**). We also noted a slight but significant reduction in the number of taste buds in the circumvallate papilla of heterozygous mice compared to wild-type control, suggesting a gene dosage effect (**Fig. 1C**).

It has been well documented that including the *Neo* cassette in an intron of a floxed gene interferes with the expression of that gene (23). Would this be the case for *Rspo2^Neo/Neo^* mice, in which *Neo* cassette insertion creates a hypomorphic allele? To address this, we performed quantitative reverse-transcription PCR (qRT-PCR) analysis of Rspo2 expression in the NPJ complex. As predicted, Rspo2 expression in homozygous *Rspo2^Neo/Neo^*mice was reduced approximately 50% compared to wild-type mice (**Fig. 1D**), which well correlated with the number of taste buds in the circumvallate papilla of wild-type and homozygous *Rspo2^Neo/Neo^*mice. Thus, these data are consistent with the idea that Rspo2 is required for taste bud generation and maintenance.

### The number of fungiform papillae is significantly reduced in homozygous *Rspo2^Neo/Neo^* mice

To determine if this hypomorphic allele of *Rspo2^Neo^*also affects other taste fields, we examined fungiform papillae in the anterior tongue. Each fungiform papilla typically contains a single taste bud. Because fungiform papillae scatter in a stereotypic pattern throughout the anterior tongue and the area surrounding the intermolar eminence, we used scanning electron microscopy to identify fungiform papillae in the dorsal surface of tongue of wild-type, heterozygous, and homozygous *Rspo2^Neo/Neo^* mice (**Fig. 2A**). Similar to what we found in the circumvallate papilla, many fewer fungiform papillae were present in homozygous *Rspo2^Neo/Neo^* mice than in wild-type or heterozygous mice (**Fig. 2B**). Nevertheless, the morphology of fungiform papillae visualized from the dorsal surface or taste buds in sections appeared to be comparable (**Fig. 2C**). Furthermore, we observed no apparent differences in Krt8 or P2x3 immunostaining in fungiform taste buds (**Fig. 2D**). Whole-mount immunostaining of peeled lingual epithelium confirmed that there are fewer taste buds in homozygous *Rspo2^Neo/Neo^* mice than in wild-type mice (**Fig. S3**).

**Fig. 2.**
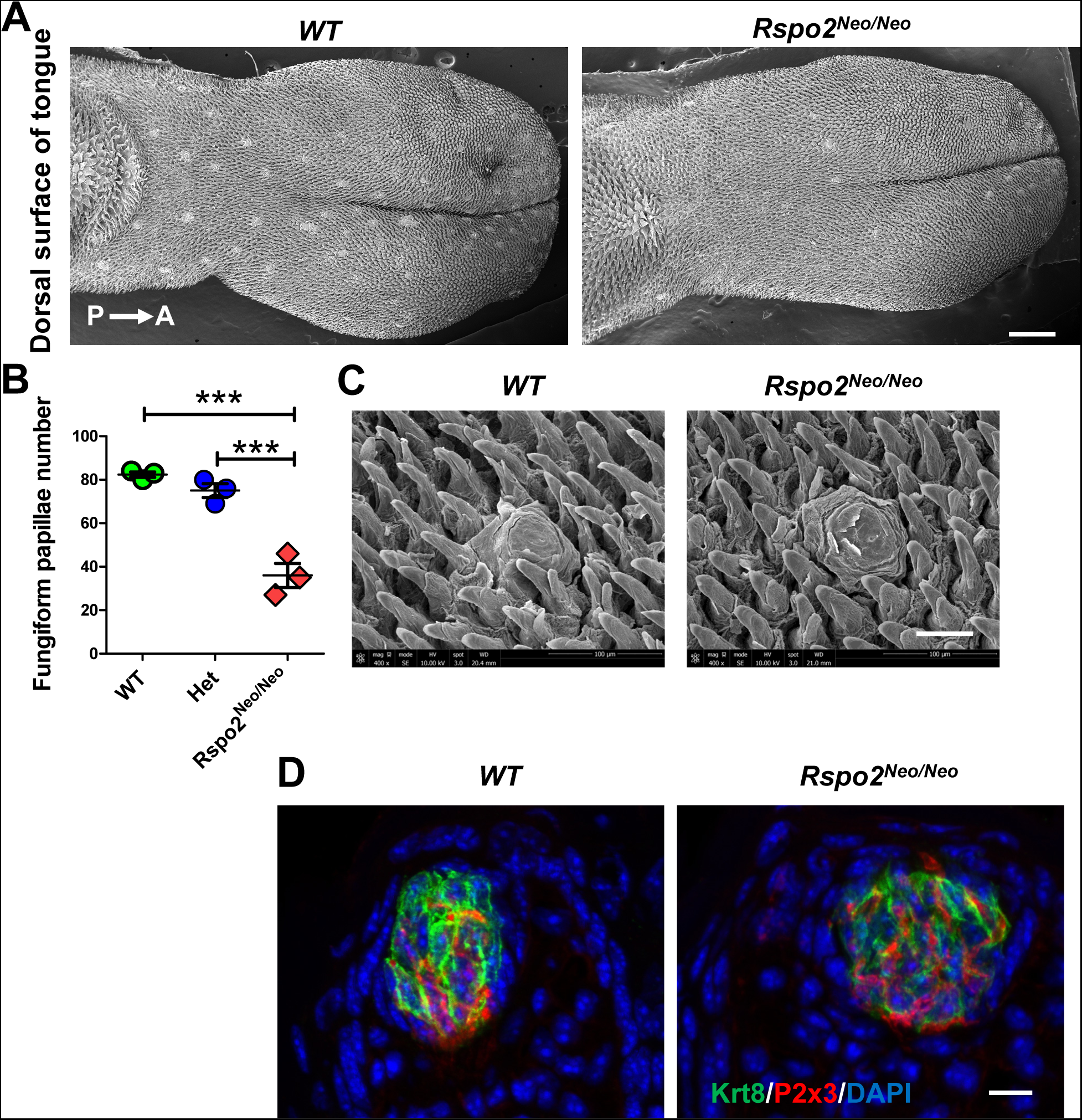
Fewer fungiform taste buds in the anterior tongue of *Rspo2^Neo/Neo^* mice than in wild-type mice. **A)** Representative scanning electron microscopy images of the dorsal surface of tongues of wild-type (WT) and *Rspo2^Neo/Neo^* mice. Fungiform papillae (whitish spots) are scattered in a stereotypic way in wild-type mice, and fewer fungiform papillae are present in *Rspo2^Neo/Neo^*mice. Orientation is from posterior to anterior (P→A). Scale bar: 0.5 mm. **B)** Number of fungiform papillae in the dorsal surface of the tongue (from tip to the intermolar eminence). Note that fungiform papillae in the tip of the ventral surface were not visible and thus were not counted. Significant differences were noted between *Rspo2^Neo/Neo^* mice and wild-type or heterozygous mice (***p <0.001). Each point represents a single mouse. Three mice were used for each genotype. **C)** Representative fungiform papillae of wild-type (WT) and *Rspo2^Neo/Neo^*mice at higher magnification. Scale bar: 50 μm. **D)** Representative confocal images of taste buds immunostained with Krt8 (green) and P2x3 (red) in fungiform papillae of wild-type (WT) and *Rspo2^Neo/Neo^* mice. Scale bar: 10 μm.

Despite a reduced number of taste buds in *Rspo2^Neo/Neo^* mice compared to wild-type mice, there was no detectable change in the number of neurons (indicated by labeling with the AM1-43 fluorescent probe) or the percentage of P2x3+ neurons in either the geniculate ganglion or NPJ complex between homozygous *Rspo2^Neo/Neo^* and wild-type mice (**Fig. S4**). These data therefore exclude the possibility that the reduced number of taste buds is caused by changes in the number of ganglion neurons that innervate taste tissues in homozygous *Rspo2^Neo/Neo^*mice. Instead, these data support a role of endogenous Rspo2 in the generation and/or maintenance of both fungiform and circumvallate papilla taste buds.

### Removal of the *NeoR* gene restores the number of taste buds

To confirm that inclusion of the *Neo* cassette was responsible for the Rspo2 hypomorphic allele, we excised the *Neo* cassette that is flanked by two FRT sites by crossing with a transgenic strain (24) expressing FLPe recombinase (**Fig. 3A**). Deletion of the *Neo* cassette in the floxed Rspo2 mice (termed *Rspo2^ΔNeo/ΔNeo^*) restored to normal the number of taste buds in the circumvallate papilla (**Fig 3B,C**). Thus, these data provide further support that Rspo2 is essential for taste bud generation and maintenance.

**Fig. 3.**
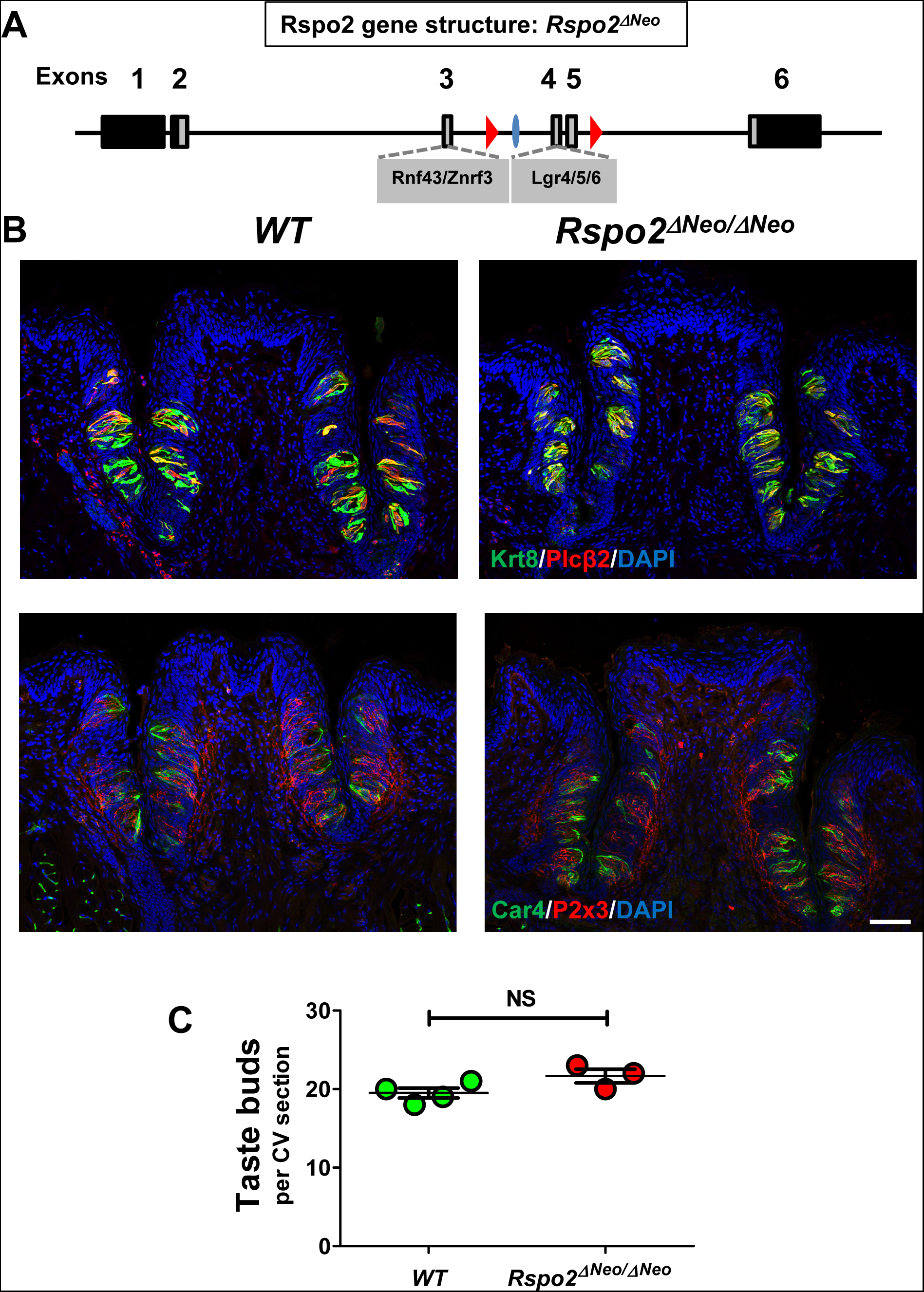
The number of taste buds returns to normal after genetic removal of the *Neo* cassette in *Rspo2^Neo/Neo^*mice. **A)** Schematic drawing showing the genetic structure of the *Rspo2* allele after removal of the *Neo* cassette (compare to Fig. 1A), leaving one FRT site (blue oval). Red triangles depict LoxP sites. **B)** Representative confocal images of circumvallate papilla sections of wild-type (WT) and *Rspo2^ΔNeo/ΔNeo^* mice immunostained with Krt8 (top row; green) and Plcβ2 (red) or Car4 (bottom row; green) and P2x3 (red). Scale bar: 50 μm. **C)** Statistical analysis of taste buds with Krt8+ and Plcβ2+ cells in the circumvallate papilla of wild-type (WT) and *Rspo2^ΔNeo/ΔNeo^* mice. No significant difference was found between wild-type and *Rspo2^ΔNeo/ΔNeo^* mice. Each point represents a single mouse. Three mice were used for each genotype.

### Ablation of Rspo2 specifically in the nodose-petrosal-jugular ganglion complex leads to loss of taste buds in the circumvallate papilla

We demonstrated that Rspo2 is required for the generation and maintenance of taste buds: many fewer taste buds are present in *Rspo2^Neo/Neo^*mice than in wild-type mice, owing to expression of Rspo2 at a reduced level, both constitutively and globally. We previously showed that during adulthood Rspo2 is expressed in gustatory neurons, not in taste tissue or in surrounding tissues (19). However, we cannot completely rule out the possibility that potential non-neuronal sources of Rspo2, during early development or beyond, may also contribute to fewer taste buds in *Rspo2^Neo/Neo^* mice than in wild-type control mice.

Furthermore, the original strategy for tissue-specific knockout of Rspo2 (see Fig. 1A) in *Rspo2^Neo/Neo^*and *Rspo2^ΔNeo/ΔNeo^* mice was to conditionally ablate exons 4 and 5 that encode the Lgr4/5/6 binding domain (25). Our recent work showed that ablation of Rnf43/Znrf3, two other receptors for R-spondin, mirrors the effect of exogenous R-spondin on taste tissue generation and regeneration (19, 20). Rspo2 interacts with Rnf43/Znrf3 via a Furin-1ike domain encoded by exon 3 (26). Because all coding exons are in-frame, the Cre-based deletion strategy may leave Rnf43/Znrf3 binding intact in *Rspo2^Neo/Neo^*and *Rspo2^ΔNeo/ΔNeo^* mice.

To circumvent such a limitation, we acquired another strain with insertion of two LoxP sites flanking exons 3-6 of Rspo2, which would permit ablation of both the Rnf43/Znrf3 and the Lgr4/5/6 binding domains using the Cre-based approach. Nevertheless, the two LoxP sites are far apart from each other (>70 kbp), and a strong Cre drive might be warranted for specific deletion of Rspo2 using this new strain (termed *Rspo2^Gem-fl/Gem-fl^*) (**Fig. 4A**).

**Fig. 4.**
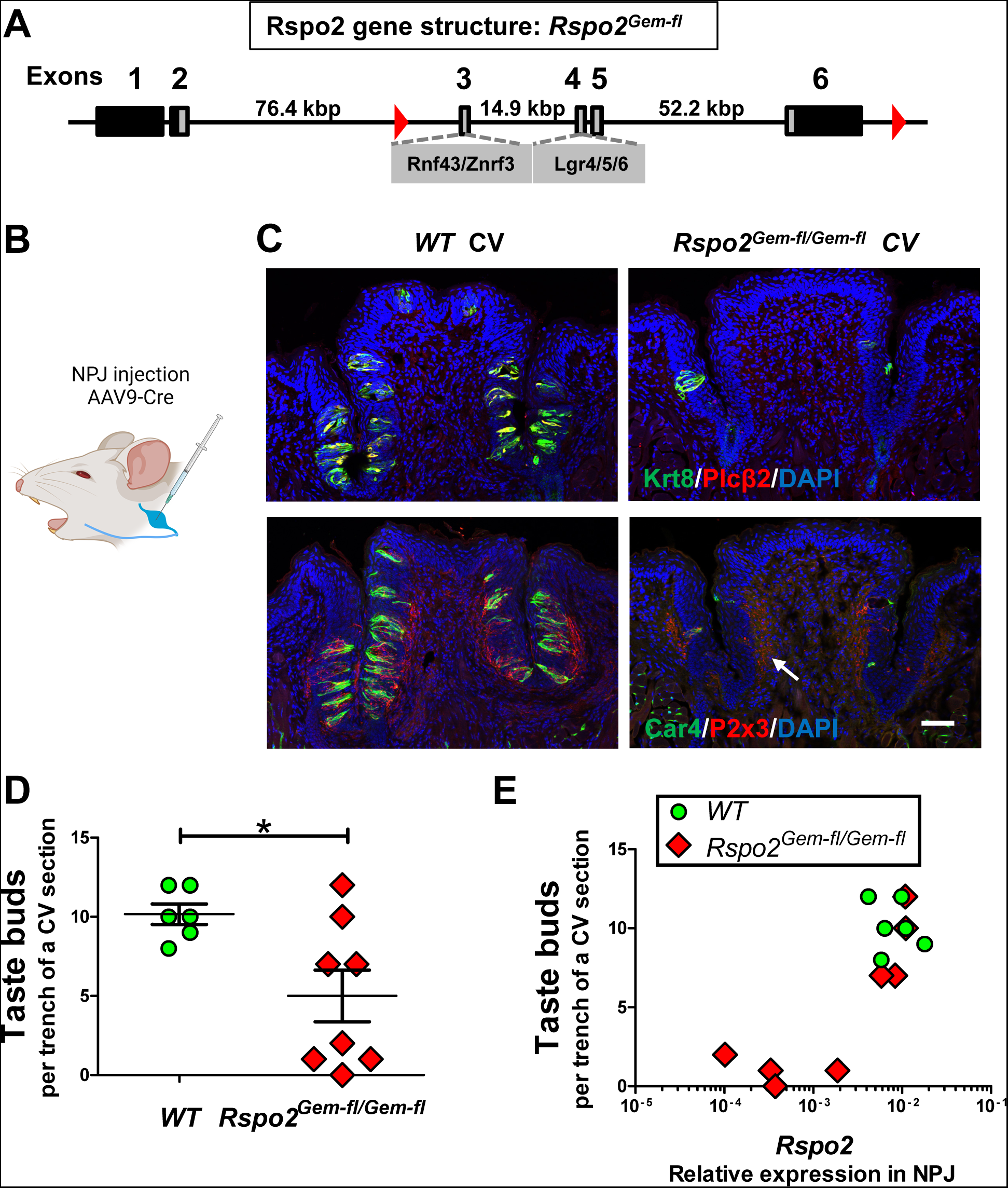
NPJ-specific deletion of Rspo2 leads to a nearly complete loss of taste buds in the circumvallate papilla. **A)** Schematic drawing illustrates the insertion of two LoxP sites (red triangles) in the *Rspo2* gene locus to remove exons 3-6 encoding the binding domains for both Lgr4/5/6 and Rnf43/Znrf3 in a Cre-dependent manner in *Rspo2^Gem-fl/Gem-fl^* mice. **B)** Schematic illustration showing the viral-based ablation of Rspo2 in NPJ (generated using BioRender). **C)** Confocal images of circumvallate papilla (CV) sections from wild-type (WT) and *Rspo2^Gem-fl/Gem-fl^* mice injected with AAV9-Cre bilaterally in the NPJ complex a month prior to tissue collection, immunostained with antibodies against Krt8 (top; green) and Plcβ2 (red) or Car4 (bottom; green) and P2x3 (red). Note the nearly complete loss of taste buds and retraction of P2x3+ fiber terminals (arrow), which sit right underneath the epithelium, after AAV9-Cre induced ablation of Rspo2 in NPJ. Scale bar: 50 μm. **D)** Number of taste buds that comprise Krt8+ and Plcβ2+ cells in the two (left and right) trenches of circumvallate papilla (CV) of wild-type (WT) and *Rspo2^Gem-fl/Gem-fl^* mice after AAV9-Cre-induced ablation of Rspo2 in NPJ. Significant difference was found between wild-type and *Rspo2^Gem-fl/Gem-fl^* mice (*p <0.05). Each point represents a single trench region from a mouse. Three to four mice were used for each genotype. **E)** Number of taste buds in each trench of the circumvallate papilla (CV) of wild-type (WT) and *Rspo2^Gem-fl/Gem-fl^* mice inoculated with AAV9-Cre against the expression level of Rspo2 in the corresponding NPJ (two NPJ complexes per mouse). Note that effective deletion of Rspo2 in an NPJ complex corresponds to nearly complete loss of taste buds in the trench area innervated by that NPJ.

*Rspo2^Gem-fl/Gem-fl^* mice have a normal number of taste buds in the circumvallate papilla (**Fig. S5**). Because we were unable to identify a CreERT2 strain that is adequate for ablating Rspo2 in gustatory neurons, we decided to ablate Rspo2 using a viral-based strategy. AAV-mediated delivery of Cre has been used for precise targeting of genes in the nervous system. Therefore, we tested if microinjection of AAV-hSyn-Cre-P2A-dTomato (hereafter referred to as AAV9-Cre) into the NPJ complex (**Fig. 4B**) would lead to deletion of Rspo2 in gustatory neurons and subsequent loss of taste buds in the circumvallate papilla, which is innervated by petrosal ganglion neurons.

Initially, we performed bilateral injection of AAV9-Cre into NPJ of *Rspo2^Gem-fl/Gem-fl^* mice and examined the circumvallate papilla a month later, to allow sufficient time for taste cell turnover and taste bud degeneration, to determine if neuron-produced R-spondin had induced and maintained taste tissue homeostasis. In two out of three mice we noted a nearly complete loss of taste buds and retraction of P2x3+ fiber terminals in the lateral walls surrounding the trench, as expected, on either one or both sides of the circumvallate papilla (**Fig. S6**). The one mouse with little change in the number of taste buds in the circumvallate papilla likely arose from technical challenges in delivery of a small amount of virus to the NPJ complex, and consequent variation in efficiency of ablation of Rspo2 from NPJ neurons.

To further explore this, we repeated this experiment using additional mice but also included wild-type mice as controls and collected each NPJ complex (left and right separately), as they innervate the corresponding left and right trenches of the circumvallate papilla. We used qRT-PCR analysis of Rspo2 expression to determine if there is a match between Rspo2 expression and the number of taste buds in the lateral wall of each trench.

Injection of AAV9-Cre in the NPJ complex of control wild-type mice did not change the number of taste buds or the innervation pattern (**Fig. 4C, left**). Similar to what we described above, in three out of four *Rspo2^Gem-fl/Gem-fl^*mice we noted a nearly complete loss of taste buds (**Fig. 4C, right**) and retraction of P2x3+ fiber terminals (**Fig. 4C, lower right**) in one or both trenches, a pattern that differs from that in mice subject to glossopharyngeal nerve transection (19). Plotting the number of taste buds in each trench area revealed a significant reduction in the number of taste buds in homozygous *Rspo2^Gem-fl/Gem-fl^* mice (**Fig. 4D**) compared to wild-type control mice, despite variability among the homozygous mice. When we compared the expression level of Rspo2 via qRT-PCR in the NPJ complex and the number of taste buds in the lateral walls of the corresponding trench, we found a nearly perfect match: a significant reduction in the expression of Rspo2 in NPJ by AAV-Cre-mediated ablation of the *Rspo2* gene (exons 3-6) led to a nearly complete loss of taste buds in the corresponding area in the circumvallate papilla (**Fig. 4E**). These data clearly demonstrate that Rspo2 produced by gustatory neurons regulates taste tissue homeostasis.

## DISCUSSION

Gustatory neurons induce and maintain taste tissue homeostasis in mammals. Our recent work suggested that Rspo2 is the long-sought gustatory-neuron-produced factor that regulates taste stem/progenitor cell activity for taste tissue maintenance (19, 20). Until now, the necessity of Rspo2 has not been demonstrated *in vivo*. In the present work, we show that a hypomorphic allele of *Rspo2* leads to a substantially reduced number of taste buds in both the anterior and posterior tongue in mice. Additionally, we show that NPJ-complex-specific ablation of Rspo2 can lead to a nearly complete loss of taste buds. These data, along with our previous work, provide compelling evidence to support the case that Rspo2 is the long-sought neuronal factor that induces taste bud formation and regulates taste tissue homeostasis.

In the strain that carries the Rspo2 hypomorphic allele, the expression of Rspo2 in NPJ is about half of the amount found in NPJ of wild-type mice, due to the insertion of *NeoR* in the intron region of *Rspo2*. The reduced Rspo2 expression correlates with a reduced number of taste buds in both anterior and posterior tongue. The removal of *NeoR* restores the number of taste buds in the circumvallate papilla. Interestingly, taste buds in *Rspo2^Neo/Neo^* mice appear to be normal, with the presence of taste receptor cells (e.g., type 2 Plcβ2+ cells and type 3 Car4+ cells), and are densely innervated with P2x3+ nerve fibers, with no noticeable change in the number of gustatory neurons.

Why does reduced expression of Rspo2 in gustatory neurons lead to fewer taste buds instead of much smaller taste buds? A stereotypic connection has been suggested between taste buds and gustatory neurons: one taste bud (e.g., in fungiform papilla or soft palate) is typically innervated by three to five ganglion neurons that primarily connect with that bud (27). Although this is perhaps a simplified view (28), on average there is a linear (1:4) relationship between taste buds injected with a lipophilic dye and the number of gustatory neurons labeled by such dye (27). Theoretically, these three to five gustatory neurons may provide the amount of Rspo2 capable of activating Lgr4/5/6 and/or silencing Rnf43/Znrf3 (receptor-ligand interaction) to maintain taste bud homeostasis. In the case of *Rspo2^Neo/Neo^* mice with the hypomorphic *Rspo2* allele, it is plausible that more neurons in *Rspo2^Neo/Neo^* mice than in wild-type mice converge to provide the amount of Rspo2 required to induce single taste buds during taste bud ontogeny, as the amount of Rspo2 produced in gustatory neurons in *Rspo2^Neo/Neo^* mice may only be half of that in wild-type mice. Therefore, fewer taste buds will be induced and maintained in the oral cavity in *Rspo2^Neo/Neo^* mice than in wild-type mice, as there is no discernible structural change in gustatory ganglia between *Rspo2^Neo/Neo^*and wild-type mice, and both have comparable numbers of gustatory neurons. This warrants further exploration in the future to determine the innervation pattern of taste buds using this model.

Consistent with the phenotypic change we noted with the strain carrying the Rspo2 hypomorphic allele, we detected a nearly complete loss of taste buds in one side or both sides of the circumvallate papilla of *Rspo2^Gem-fl/Gem-fl^* mice with substantial deletion of Rspo2 (by qRT-PCR analysis) in NPJ neurons after receiving AAV9-Cre injection. When Rspo2 was not substantially deleted, presumably due to technical challenges in microinjection or other causes, the number of taste buds did not change much. Amazingly, the level of Rspo2 in an NPJ complex matches almost perfectly the number of taste buds in the corresponding side of the circumvallate papilla, in line with the ipsilateral projection pattern of geniculate ganglion neurons. Again, this clearly establishes the necessity of Rspo2 for maintaining taste tissue homeostasis and provides a compelling explanation for the long-ago observation that taste bud maintenance requires innervation.

In concordance with this, we previously showed that systemic provision of R-spondin substitutes for neuronal input for generation of taste buds and that epithelial ablation of Rnf43/Znrf3 phenocopied the effect of exogenous R-spondin (19, 20). Interestingly, we observed that ectopic taste buds were generated in mice receiving exogenous R-spondin or epithelial ablation of Rnf43/Znrf3 (19, 20). How these buds were generated remains unclear. One possibility is that exogenous R-spondin or ablation of Rnf43/Znrf3 (the negative regulator) may activate quiescent taste stem cells that are pre-fate-determined either to be quiescent or to generate non-taste epithelial cells in the absence of innervation. Another possibility is that exogenous R-spondin or ablation of Rnf43/Znrf3 will expand taste stem/progenitor cells (e.g., Lgr5+ basal cells in posterior tongue) to produce more taste buds. Regardless, during normal homeostasis gustatory innervation dictates taste bud patterning via Rspo2.

Despite our demonstration of the necessity and sufficiency of Rspo2 for neuronal regulation of taste stem cells, it remains unclear how Rspo2 is released from gustatory neurons. It would be useful to know the mechanism underlying the secretion of Rspo2. Perhaps a mechanism somewhat similar to secretion of neurotropic factors such as BDNF, NGF, or NT3 is utilized for secretion of R-spondin (29). This remains to be addressed.

R-spondin is expressed not only in gustatory neurons but presumably also in vagal neurons and beyond. For instance, our previous *in situ* hybridization work showed that R-spondin transcript was detected across the NPJ complex, which suggests that certain nodosal neurons may produce Rspo2 as well (19). Nevertheless, the effects we observed in the circumvallate papilla are more likely mediated by gustatory neurons of the petrosal ganglion because they uniquely innervate taste buds. Moreover, recent single-cell RNA sequencing reveals that Rspo2 is expressed in certain dorsal root ganglion neurons (e.g., those expressing Mas-related G protein-coupled receptors [Mrgprs]) and enteric neurons (30). The function of Rspo2 in those neurons remains to be characterized. Nevertheless, it has been reported that Lgr6 epidermal stem cells are primed for injury response in a nerve-dependent manner (31). Perhaps, Rspo2 (and/or Rspo1) produced by the dorsal root ganglion neuron (30) is involved in this process.

Furthermore, enteric neurons express Rspo2 (30). The role of Rspo2 expressed in vagal and enteric neurons remains unclear. These neurons may contribute to intestinal stem cell homeostasis. Recent data suggests that Rspo3 from lymphatic endothelial cells and RSPO3^+^GREM1^+^ fibroblasts contribute to such a process (32), but additional regulation from enteric neurons or other neurons may be a key component as well.

Regardless, taste innervation is one of the best examples to reveal the central role of neuronal Rspo2 in regulating the homeostatic maintenance of distant sensory organs.

## METHODS

### Mice

All experiments were performed under National Institutes of Health Guidelines for the Care and Use of Animals in Research and approved by the Institutional Animal Care and Use Committee of the Monell Chemical Senses Center. The *Rspo2^Neo/Neo^* strain was provided by Dr. Kurt Hankenson under a materials transfer agreement. The *Rspo2^Gem-fl/Gem-fl^*strain (T011127) was acquired from GemPharmatech. All other strains (e.g., B6.Cg-Tg(ACTFLPe)9205Dym/J, Tg(Thy1-cre/ERT2,-EYFP)HGfng/PyngJ) were obtained from Jackson Laboratory (JAX #005703, #012708, respectively). All genetically modified mice were maintained on the C57BL/6 genetic background. Both adult males and females between 8 and 18 weeks old were used. Littermates or age-matched mice with the same genetic background (C57BL/6J) were used throughout the study.

### AAV injection into the NPJ complex

The mice were given an injection of the NSAID carprofen (5 mg/kg) and underwent anesthesia (isoflurane, 1.5-2.5%). They were then placed in a supine position on a heating pad, and a 2 cm midline incision was made in the skin on the ventral side of the neck. After moving aside the skin, salivary glands, and underlying muscles, the vagus nerve was exposed using fine-tip forceps. The NPJ complex was located by tracing the vagus nerve toward the base of the skull and was then exposed by retracting adjacent muscles and blunt dissection of connective tissues. A glass micropipette filled with the viral construct (pAAV-hSyn-Cre-P2A-dTomato; Addgene, #107738-AAV9; titer ≥ 1 × 10^13^ viral genomes/mL) attached to a micromanipulator was used to position and puncture the NPJ, with ~0.5 µl injected bilaterally into the NPJ using a Picospritzer III injector (Parker Hannifin, Pine Brook, NJ). The incision was closed, and after 24 hr post-op analgesic was administered. Tissues were harvested 1 month later (between 31-33 days).

### Immunostaining and imaging

Immunostaining was performed essentially as described previously (19, 20). When only tongues were collected, mice were not perfused; when ganglia were collected for immunohistochemistry, mice were perfused. Tongues and ganglia were fixed in 4% (wt/vol) paraformaldehyde (PFA) for 2 hr, cryoprotected with 30% sucrose overnight, embedded in OCT, and sectioned at either 10 μm (taste tissue, coronal sections for the circumvallate papilla, and sagittal sections for the anterior part of the tongue/fungiform papillae) or 16 μm (ganglia, horizontal sections). To identify taste cells, we used specific primary antibodies against taste cell markers and species-specific secondary antibodies: goat anti-Car4 (1:100; R&D, #AF2414), rabbit anti-P2x3 (1:1000; Alomone Labs, APR-016), rabbit anti-Plcβ2 (1:500; Santa Cruz, #SC-206), and rat anti-Krt8 (1:10; Developmental Studies Hybridoma Bank, #Troma-1). For sections from wild-type mice double staining was performed, such as pairs of anti-Car4 and anti-P2x3 antibodies, or anti-Plcβ2 and anti-Krt8 antibodies, or anti-P2x3 and anti-Krt8 antibodies. To identify P2x3+ neurons, geniculate and NPJ sections were stained with the antibody against P2x3. Fluorescence-labeled secondary antibodies against specific species were used to visualize staining. All images were acquired by a Leica DMi8 confocal microscope, or a Leica TCS SP8 confocal microscope at the Cell and Developmental Biology Core at the University of Pennsylvania, or a Nikon Eclipse E800 or Olympus BX63 microscope. Confocal images were compressed z-stacks of the entire section (~10 μm for taste tissue sections and ~16 µm for ganglion sections); single optical sections gave rise to the same results as the z-stacks.

### Taste bud counting and ganglion neuron counting

Taste buds were recognized as onion-like structures with at least three Krt8+ cells (one of which was also positive for Plcβ2 staining). The numbers of taste buds were counted manually as described previously (20). A section in the middle of the circumvallate papilla for each mouse was used for taste bud counting to alleviate potential sampling bias associated with sections at the most anterior and posterior portions of the circumvallate papilla. Ganglion neurons were counted as P2x3+ neurons when they showed clear cell-surface staining and recognizable nuclei stained with DAPI. The total number of neurons was counted by adding the number of P2x3+ neurons and P2x3 negative neurons with background fluorescence, validated using bright-field imaging. A section in the middle of NPG was used to count the P2x3+ and P2x3 negative neurons and calculate the percentage of P2x3+ neurons for each mouse. One or two sections from the geniculate ganglion were used to count neurons for each mouse. The data was tabulated for each mouse.

### Statistical analyses

Data were shown as the mean ± SEM. We used GraphPad Prism 5 software to generate graphs and perform statistical analyses. One-way ANOVA with post hoc Tukey tests was applied to determine statistical differences among three groups. Student’s t test was used when comparing two groups.

### qRT-PCR analysis

qRT-PCR analysis was performed essentially as described previously (20). Briefly, total RNA was extracted from NPJ using the PureLink RNA Micro Kit (ThermoFisher, #12183-016) plus TRIzol (Life Technologies, #15596-026). cDNA was synthesized using SuperScript™ IV VILO™ Master Mix with ezDNase™ Enzyme (Invitrogen, #11766500), and quantitative PCR was performed using Fast SYBR™ Green Master Mix Kit (Applied Biosystems, #4385612). *Gapdh* was used as control to normalize the expression levels of analyzed gene transcripts. Relative gene expression was calculated as 2^(CTTarget-CTGapdh)^. The primers used were intron spanning: 5’-GCCGCTGCTTTGATGAATGT-3’ (Rspo2 forward), 5’-TGGCCATCTTGCATCTCCTG-3’(Rspo2 reverse), 5’-TGGCCTTCCGTGTTCCTAC-3’ (Gapdh forward), 5’-GAGTTGCTGTTGAAGTCGCA-3’ (Gapdh reverse).

### Scanning electron microscopy

Scanning electron microscope experiments were carried out at the Cell and Developmental Biology Microscopy Core (Perelman School of Medicine, University of Pennsylvania). Samples were washed three times with 50 mM Na-cacodylate buffer, fixed for 2 hr with 2% glutaraldehyde in 50 mM Na-cacodylate buffer (pH 7.3), and dehydrated in a graded series of ethanol concentrations through 100% over 2.5 hr. Dehydration in 100% ethanol was done three times. Dehydrated samples were incubated for 20 min in 50% hexamethyldisilane (HMDS; Sigma-Aldrich) in ethanol followed by three changes of 100% HMDS, followed by overnight air drying as described previously (33). Then samples were mounted on stubs and sputter coated with gold palladium. Specimens were observed and photographed using a Quanta 250 FEG scanning electron microscope (FEI, Hillsboro, OR) at 10 kV accelerating voltage.

### Whole-mount imaging of lingual epithelium

Mice were euthanized with CO_2_ and cervical dislocation. The tongue was immediately removed and transferred into a 35-mm dish containing 4°C Hanks’ balanced salt solution (HBSS). Then, 0.5 ml of a mixture of 1 mg/ml elastase, 2 mg/ml dispase II, and 1 mg/ml trypsin inhibitor dissolved in Ca^2+^/Mg^2+^-free HBSS (HBSS(-)) was injected below the tongue epithelia, from the posterior to the anterior end of the tongue, with a 27G insulin syringe. The tissue was incubated in HBSS(-) for 30 min at room temperature (RT) and then immediately fixed in 4% PFA (Electron Microscopy Sciences, #15710) in PBS for 1 hr at RT. The peeled tongue epithelial sheet was transferred into a 15-ml tube, rinsed with PBS for 10 min at RT three times, and blocked by Superblock including 0.3% Triton X-100 and 2% donkey serum for 1 hr; immunostaining was then performed using the anti-Krt8 antibody.

### Tissue clearing of the geniculate and nodose-petrosal-jugular ganglia

The mouse received intraperitoneal administration of AM1-43 (Biotium, #70024) dissolved in PBS (2 mg/kg). After one day, the left and right geniculate ganglia and NPJ complex were isolated after cardiac perfusion with 4% w/v PFA in phosphate-buffered saline. The isolated ganglia underwent a dehydration process with ethanol, progressing through concentration steps of 50%, 75%, and three consecutive immersions in 100%, with each step taking 30 min (2.5 hr total). Subsequently, the dehydrated tissue was immersed in a BABB solution (benzyl alcohol and benzyl benzoate in a 1:2 ratio; Sigma Aldrich). The cleared ganglia were subsequently mounted between a slide glass and a coverslip. Volumetric images of the cleared ganglia were acquired with a two-photon laser-scanning microscope with an excitation wavelength tuned to 920 nm (Ultima 2P, Bruker). The images were z-projected, and the area of the ganglion was quantified by using ImageJ.

## ACKNOWLEDGMENT

We thank all members of the Jiang laboratory and Dr. Karen Yee for their input. Research reported in this publication was supported by NIH grants DC018627 and DC010842 (to P.J.) and DC015491 (to I.M.). Imaging was performed at the Cell and Developmental Biology Microscopy Core at the University of Pennsylvania and at the Monell Histology and Cellular Localization Core, which is supported in part by NIH–National Institute on Deafness and Other Communication Disorders core grant DC011735 (to R.F.M.).

## Author contributions

J.X., A.A., and P.J. designed experiments; J.X., A.A., R.X., X.L., C.L., M.Z., and M.C. performed experiments; J.X., A.A., R.X., M.C., and P.J. analyzed data; J.X., A.A., R.X., R.F.M., I.M., M.C., G.d.L., and P.J. interpreted results of experiments; K.H. contributed reagents; P.J. conceived of the work and wrote the manuscript with J.X., G.d.L., A.A., and M.C.

---

**Fig. S1.**
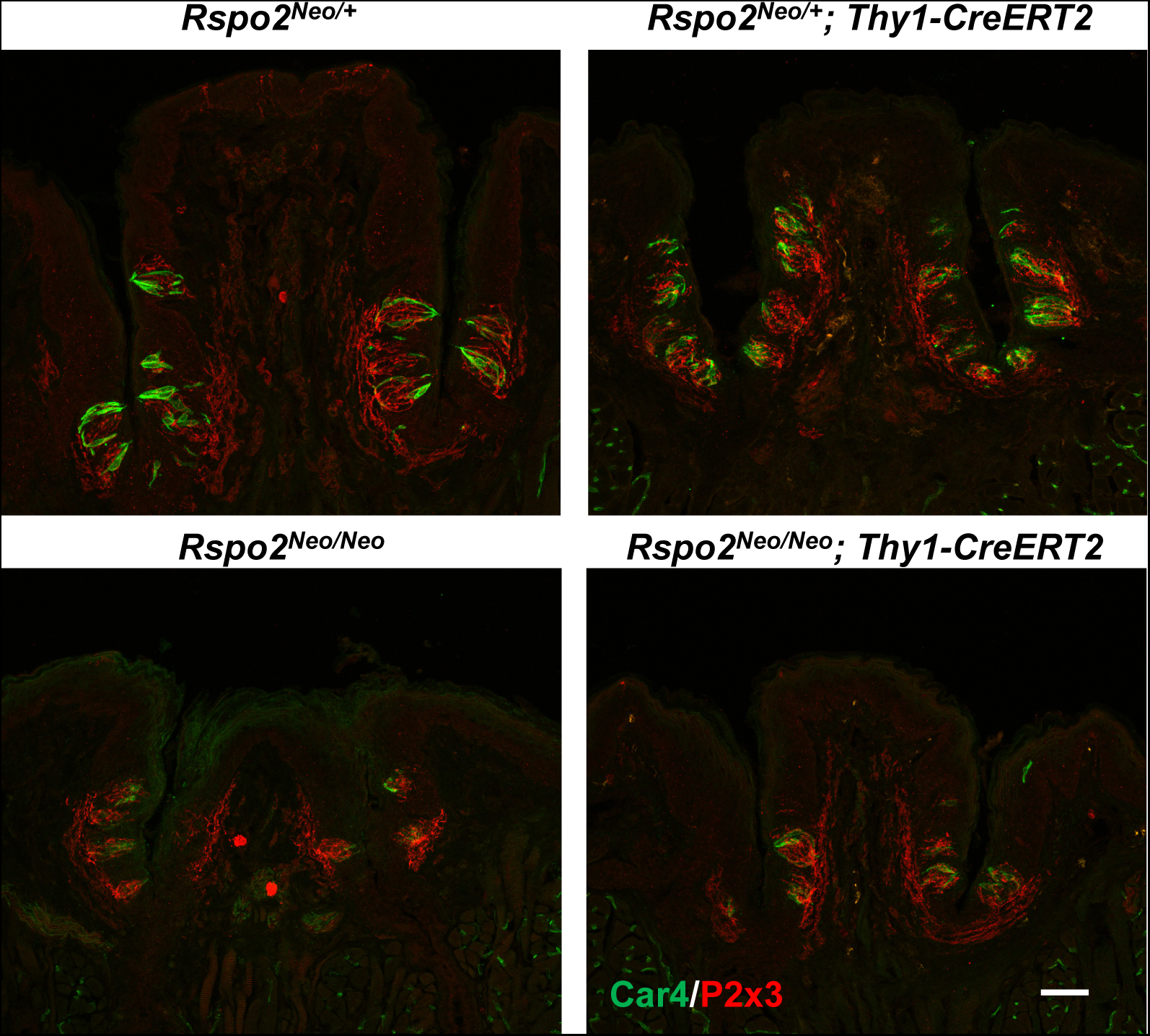
The number of taste buds in the circumvallate papilla is reduced in *Rspo2^Neo/Neo^ mice* regardless of the presence or absence of the *Thy1-CreERT2, -EYFP* transgene. *Rspo2^Neo/Neo^* mice (bottom row) in the presence (right) or absence (left) of the *Thy1-CreERT2, -EYFP* transgene were treated with tamoxifen daily for 5 consecutive days to induce Cre activity. Tissues were collected 28 days later. Circumvallate papilla sections from different genotypes were immunostained for Car4 (green) and P2x3 (red). Representative confocal images are shown. There is no indication that the presence of *Thy1-CreERT2, -EYFP* affects the number of taste buds in either *Rspo2^Neo/Neo^* or *Rspo2^Neo/+^* (top row) mice after tamoxifen induction. Scale bar: 50 μm.

**Fig. S2.**
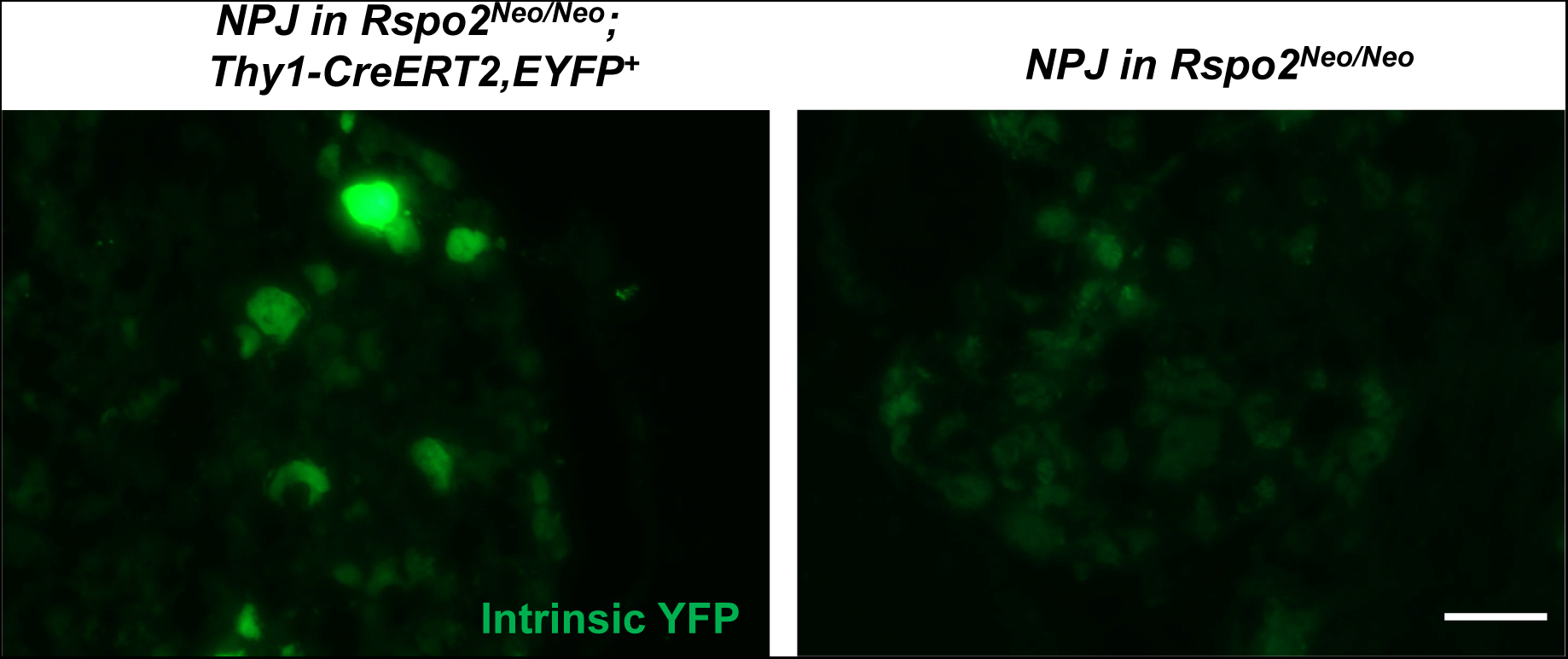
Only a limited number of neurons in the NPJ complex express the *Thy1-CreERT2, -EYFP* transgene. These representative images show only a few neurons displaying intrinsic enhanced YFP fluorescence in NPJ from an *Rspo2^Neo/Neo^*; *Thy1-CreERT2, -EYFP* mouse (left) and only background signal in NPJ from an *Rspo2^Neo/Neo^*mouse (right). Scale bar: 50 μm.

**Fig. S3.**
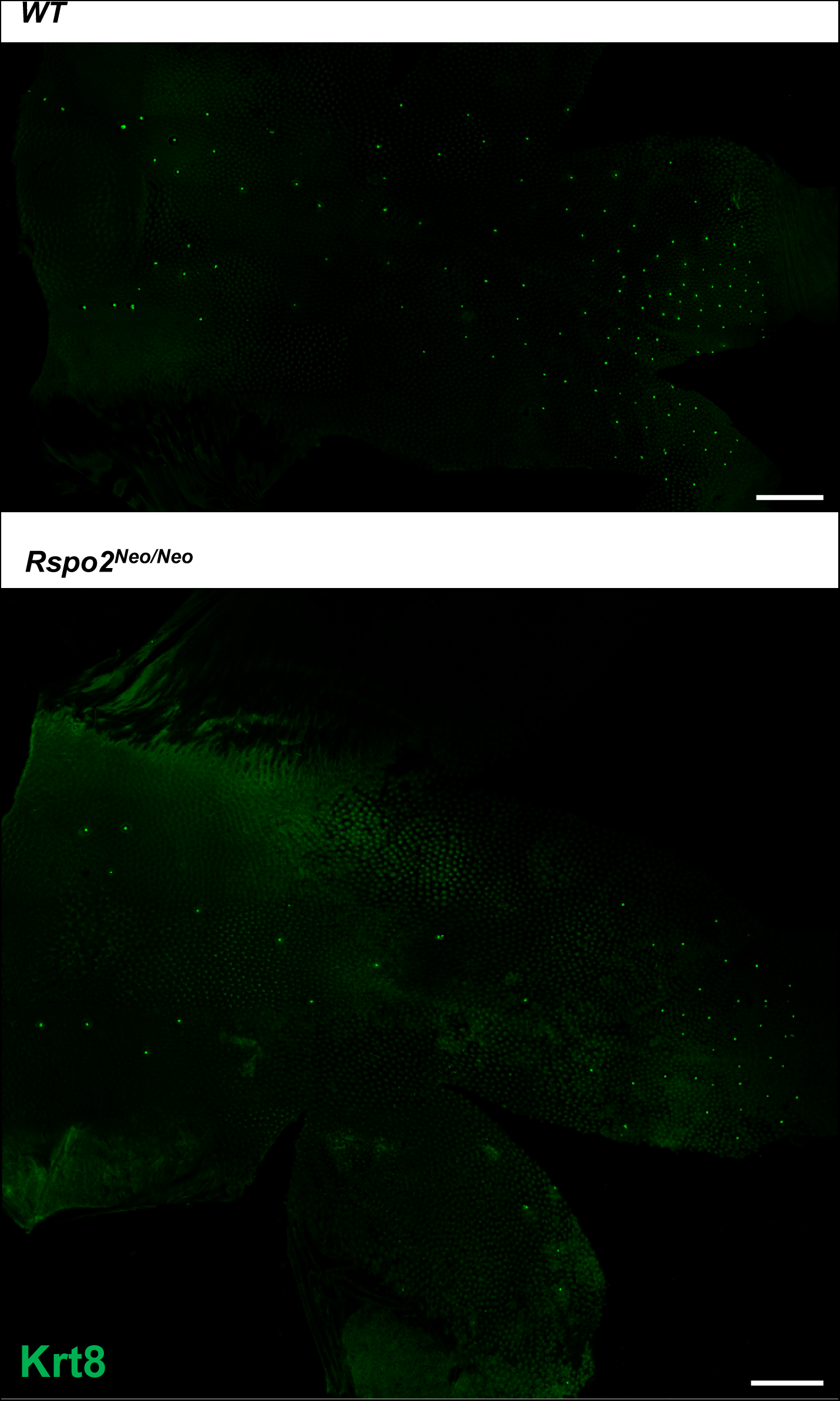
The number of fungiform papilla taste buds is reduced in *Rspo2^Neo/Neo^*mice. Whole-mount images of peeled lingual epithelium from wild-type (WT) control and *Rspo2^Neo/Neo^* mice, immunostained for Krt8 (green), show the distribution and number of fungiform taste buds. Note the fewer fungiform papilla taste buds in *Rspo2^Neo/Neo^* mice. Scale bars: 1 mm.

**Fig. S4.**
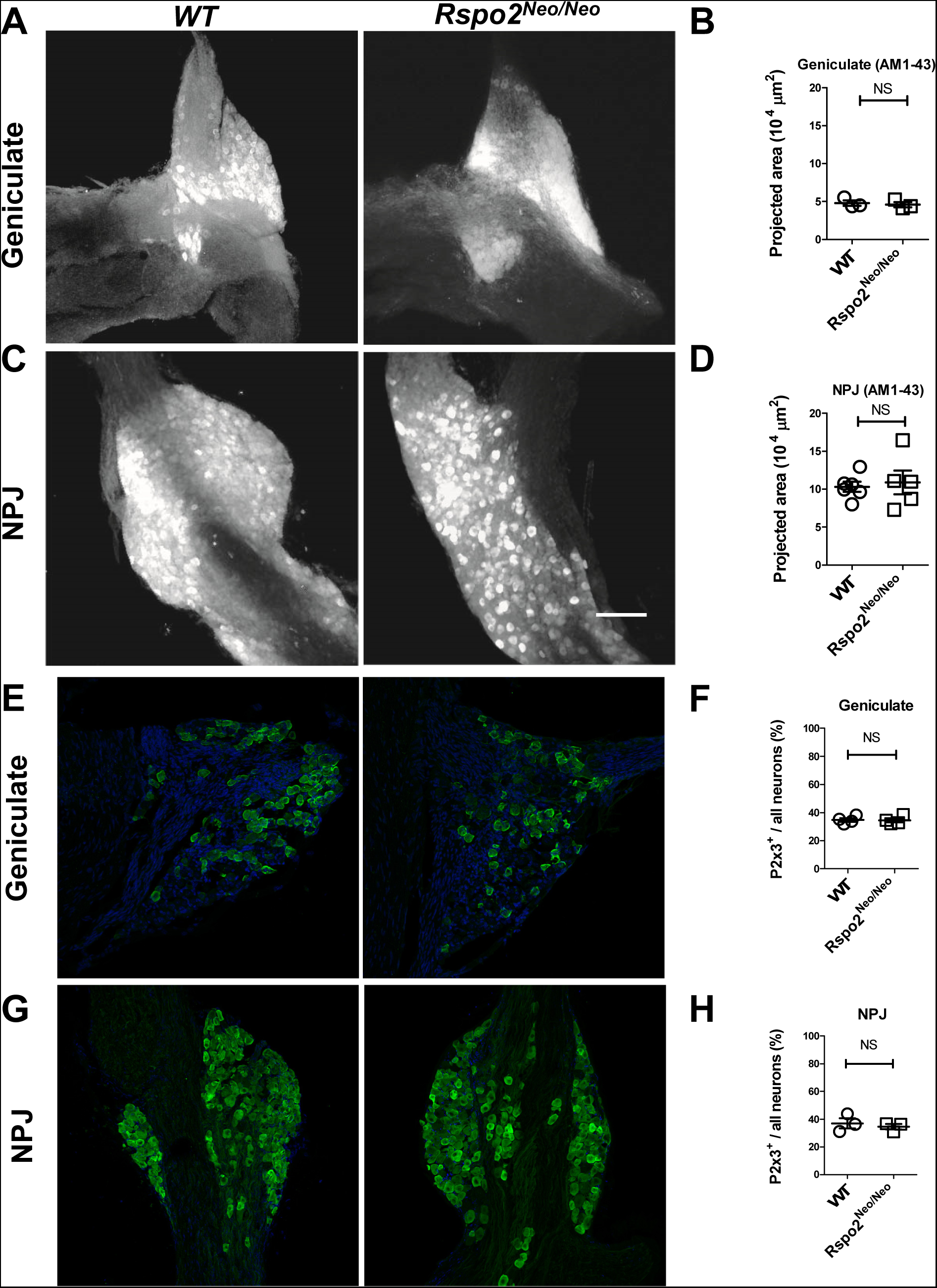
The number of neurons in the geniculate ganglion and nodose-petrosal-jugular ganglion complex appear to be comparable between wild-type and *Rspo2^Neo/Neo^* mice, so as P2x3^+^ neurons. **A)** Representative two-photon images of AM1-43-labeled geniculate ganglion from wild-type (WT) and *Rspo2^Neo/Neo^* mice. **B)** Quantification of the area comprising geniculate ganglion neurons. Each point represents a single ganglion. **C)** Representative two-photon images of AM1-43-labeled NPJ from wild-type and *Rspo2^Neo/Neo^*mice. Scale bar: 100 μm. **D)** Quantification of the area comprising NPJ neurons. Each point represents a single ganglion. **E)** Representative confocal images of P2x3+ (green) neurons in the geniculate ganglion from wild-type and *Rspo2^Neo/Neo^* mice, counterstained with DAPI. Scale bar: 50 μm. **F)** Quantification of the percentage of P2x3+ neurons over all neurons in geniculate ganglion sections. Each point represents a single mouse. **G)** Representative confocal images of P2x3+ (green) neurons in NPJ from wild-type and *Rspo2^Neo/Neo^* mice, counterstained with DAPI. Scale bar: 50 μm. **H)** Quantification of the percentage of P2x3+ neurons over all neurons in NPJ sections. Each point represents a single mouse. NS: non significant.

**Fig. S5.**
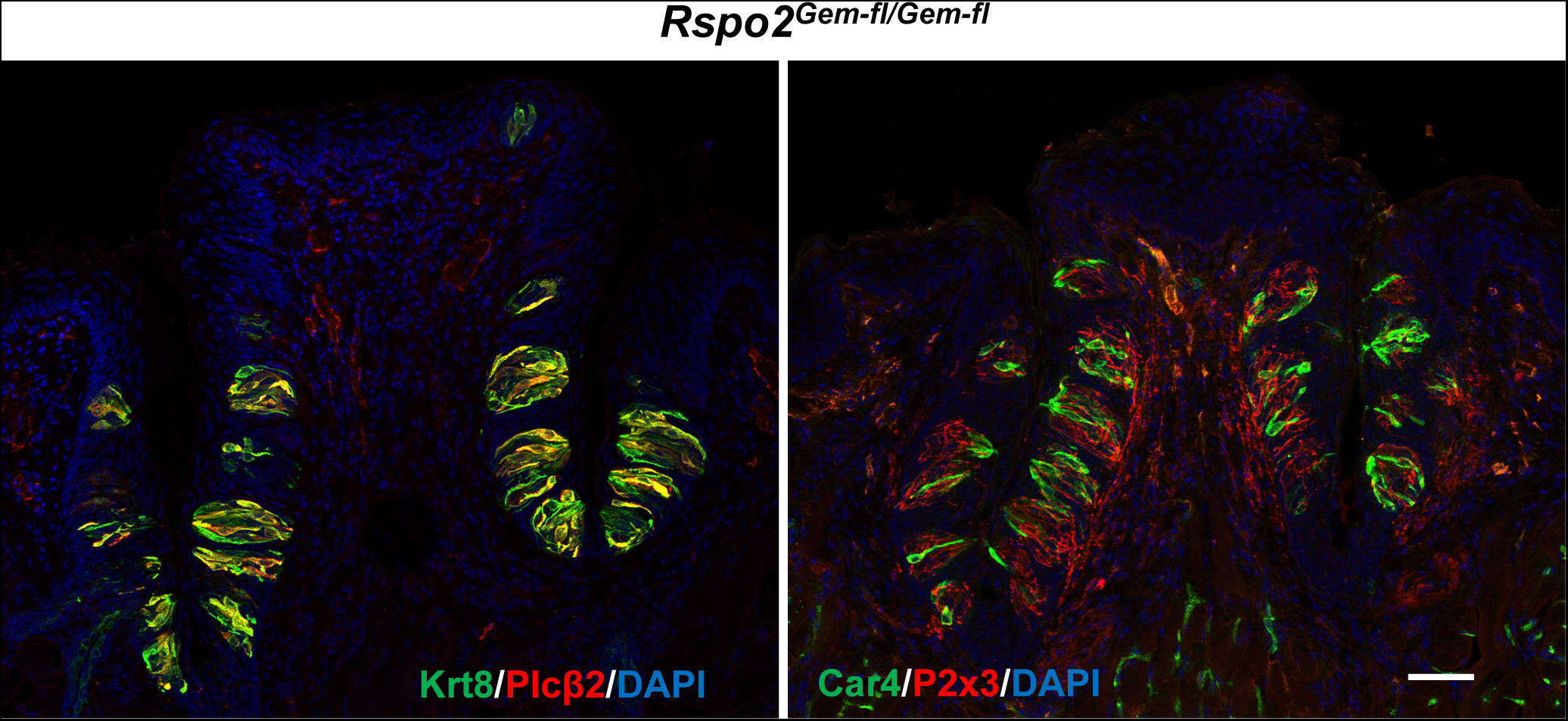
*Rspo2^Gem-fl/Gem-fl^*mice have a normal number of taste buds in the circumvallate papilla. Confocal images of immunostained circumvallate papilla sections from *Rspo2^Gem-fl/Gem-fl^*mice stained with Krt8 (left; green) and Plcβ2 (red) or with Car4 (right; green) and P2x3 (red). Scale bar: 50 μm.

**Fig. S6.**
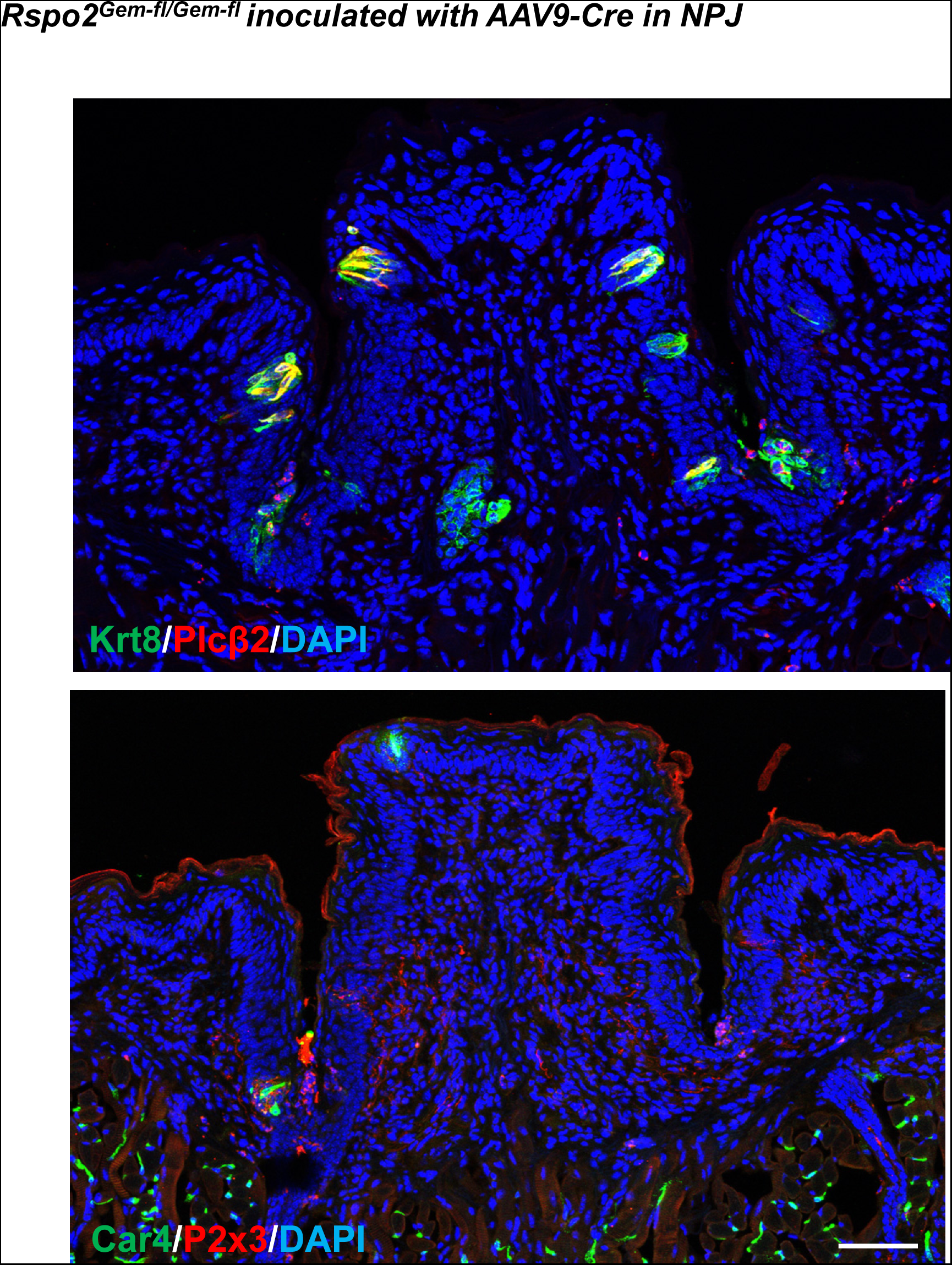
AAV9-Cre-mediated deletion of Rspo2 leads to nearly complete loss of taste buds in the circumvallate papilla. Confocal images of circumvallate papilla sections from a *Rspo2^Gem-fl/Gem-fl^*mouse (not the mouse shown in Fig. 4) injected with AAV9-Cre bilaterally in NPJ a month prior to tissue collection, immunostained with antibodies against Krt8 (top; green) and Plcβ2 (red) or Car4 (bottom; green) and P2x3 (red). Note that most taste buds degenerated after AAV9-Cre-induced ablation of Rspo2 in NPJ and that the gustatory nerves retracted. Scale bar: 50 μm.

